# Intracellular *Acinetobacter baumannii* Acts as a Reservoir in Lung Infection *via* a ‘Persist and Resist’ Strategy

**DOI:** 10.1101/2025.04.28.651116

**Authors:** Manon Janet-Maitre, Gisela Di Venanzio, Clay D. Jackson-Litteken, Nichollas E. Scott, Mario F. Feldman

## Abstract

Although considered primarily extracellular, *Acinetobacter baumannii* can survive and replicate within macrophages *in vitro*. Intracellular bacteria are often protected from the host immune system and antibiotic treatment, potentially leading to chronic or recurrent infections. To investigate the role of intracellular *A. baumannii* during infection, we transferred infected bronchoalveolar lavage fluid (BALF), containing an intracellular bacterial population, to naïve immunocompromised mice. This BALF transfer resulted in *A. baumannii* lung infection, suggesting that intracellular bacteria may serve as a reservoir for recurrent lung infections. Using dual-proteomics, we characterized the *A. baumannii*-macrophages interactions. Infected macrophages exhibit an inflammatory and Type I interferon response, marked by increased ACOD1 (IRG1) protein levels. Intracellular *A. baumannii* upregulates proteins involved in evading nutritional immunity, stress response, surface modification, and metabolic adaptation. These findings suggest that *A. baumannii* employs a multifactorial strategy to survive and replicate within macrophages, which may contribute to recurrent and chronic infections, and compromise treatment efficacy.

**Teaser:** Intracellular *A. baumannii* can act as a reservoir in lung infection

## Introduction

Hospital-acquired pneumonia burdens the USA healthcare system, raising costs, extending hospital stays, and increasing patient mortality (*1*). *Acinetobacter baumannii,* a Gram-negative opportunistic pathogen, accounts for up to 10% of hospital-acquired pneumonia in the USA and represents a major threat to global health (*2*, *3*). *A. baumannii* is also responsible for a wide variety of infections including <u>u</u>rinary tract infections (UTIs), bone and soft tissue infections and bloodstream infections (*4*, *5*). Notorious for its extremely high levels of antibiotic resistance, *A. baumannii* was classified as a critical priority pathogen for the development of new therapeutics by the <u>W</u>orld <u>H</u>ealth <u>O</u>rganization (WHO, (*6*, *7*)).

In *A. baumannii* pneumonia, <u>a</u>lveolar <u>m</u>acrophages (AMs) and lung epithelial cells are the first line of defense against bacterial infection (*4*, *8*, *9*). AMs are particularly important in the early stages of *A. baumannii* respiratory infection, where they have both direct and indirect roles. They act directly by phagocytosing *A. baumannii*, to reduce bacterial burden in the lung (*10*). In addition, they play a critical role in initiating and regulating the innate immune response, notably through the recruitment of neutrophils to the site of infection, the main cell type responsible for bacterial clearance (*10–12*). Recent studies have shown that clinical isolates of *A. baumannii* can survive and/or replicate within lung epithelial cells, endothelial cells, and both human- and mouse-derived macrophages *in vitro.* Vacuoles containing *A. baumannii* have been observed in mouse alveolar macrophages at 4- and 24-<u>h</u>ours <u>p</u>ost-infection (hpi) in an acute pneumonia model (*13*). Recently, <u>T</u>oll like receptor 4 (TLR4)-deficient mice, which are more susceptible to Gram-negative infection, were shown to develop chronic pneumonia when inoculated with low, clinically relevant bacterial dose (*14–16*). In this model, *A. baumannii* containing vacuoles (ACVs) were detected in the AMs at two days post-infection (*17*). Intracellular replication of *A. baumannii* has been mostly investigated in J774.A1 macrophages. Following phagocytosis, phagosome maturation stalls in the case of the replicative, recent clinical isolate Ab398, which survives and replicates within large, non- degradative ACVs (*13*). Notably, Ab398 counteracts phagosome acidification by secreting large amounts of ammonia, a by-product of amino acid catabolism (*13*). Intracellular replication of *A. baumannii* occurs over a short time course, peaking at 4-8h post-infection in a strain- dependent manner (*13*, *18*). In contrast, vacuoles containing the non-replicative *A. baumannii* strain ATCC19606, herein referred to as Ab19606, recruit the autophagy marker LC3, resulting in bacterial clearance. The intracellular bacterial population subsequently decreases as bacteria egress from the macrophage through a seemingly lytic process (*18*). Whether *A. baumannii* follows similar replication kinetics and characteristics *in vivo* remains unknown.

In other bacterial species, intracellular localization has been shown to protect bacteria from both the host immune response and antibiotic treatment, reducing antibiotic efficacy and creating a reservoir that can lead to chronic or recurrent infections (*19*, *20*). For example, *Escherichia coli* can form intracellular <u>b</u>acterial <u>c</u>ommunities (IBCs) or <u>q</u>uiescent intracellular reservoirs (QIRs) during UTIs, contributing to recurrent infections (*21*). *Mycobacterium tuberculosis* also exemplifies this phenomenon, as it can survive within macrophages inside granulomas for years before reactivation (*22*). Similarly, intracellular *A. baumannii* has been found to persist in bladder epithelial cells following the resolution of a UTI, leading to infection resurgence upon catheterization (*23*). However, the role of the intracellular lifestyle of *A. baumannii* during respiratory infection, the main manifestation of *Acinetobacter* infections, has not been investigated.

Intracellular pathogens use various strategies to hijack the host machinery and evade phagosome maturation (*24*). For example, *Salmonella* and *Shigella* secrete highly specialized virulence factors through the type III <u>s</u>ecretion <u>s</u>ystem (T3SS, (*25*)) or type VI <u>s</u>ecretion <u>s</u>ystems (T6SS, (*26*)). Other pathogens, like *Legionella pneumophila* or *Brucella abortus* employ a T4SS instead. *A. baumannii* lacks T3SSs and T4SSs that typically facilitate the establishment of an intracellular niche. Although most *A. baumannii* strains encode a T6SS, its expression is often silenced in clinical settings and modern clinical isolates, and all T6SS effectors identified so far target bacterial competitors (*27–29*). For intracellular *A. baumannii*, besides ammonia secretion, virulence factors influencing *A. baumannii* replication in macrophages have yet to be identified.

In this work, we aimed to investigate the *in vivo* relevance of the *A. baumannii* intracellular niche and characterize the macrophage – *A. baumannii* molecular interface. We transferred infected <u>b</u>ronchoalveolar lavage fluid (BALF), containing an intracellular *A. baumannii* population, to naive immunocompromised mice and showed that these intracellular bacteria act as a reservoir capable of seeding lung infection. We subsequently employed a comparative dual-proteomic approach to characterize the molecular interplay between intracellular *A. baumannii* and J774.A1 mouse macrophages.

## Results

### Intracellular *A. baumannii* is able to seed *de novo* infection in mouse lungs

*A. baumannii* replicates within ACVs in mouse and human macrophages *in vitro*, and it has been found within <u>p</u>oly<u>m</u>orphonuclear <u>n</u>eutrophils (PMNs) and AMs, the predominant cell types in BALF *in vivo* (*10*, *13*, *30*, *31*). However, the role and significance of intracellular *A. baumannii* during infection remain unclear. We hypothesized that intracellular *A. baumannii* could serve as a reservoir during lung infection. To test this hypothesis, we performed BALF transfer experiments from infected to uninfected mice and assessed *de novo* lung infection in the latter (*Figure 1A*). We intranasally infected TLR4-deficient C3H/HeJ mice (*32*) with ∼10^9^ CFU of either the modern clinical isolate *A. baumannii* strain Ab398, or Ab19606, a lab-adapted strain. We used C3H/HeJ mice, which are suitable for studying *A. baumannii* infection with a low inoculum, as demonstrated in a recent chronic lung infection model, and to ensure compatibility with mice used in the subsequent BALF transfer step (*17*). At 24 hpi, red blood cells-depleted BALFs were pooled and treated with colistin to eliminate extracellular bacteria. The predominant cell types present in BALF at this time point are macrophages and neutrophils (*10*). In colistin-treated BALF, these immune cells harbored an intracellular population of *A. baumannii* (*Figure 1B*). Complete elimination of extracellular bacteria was confirmed by BALF antibiotic protection assays (*Figure S1A).* Bacterial loads in lungs, kidneys and spleen were equivalent between the two strains (*Figure S1B)*.

**Figure 1.**
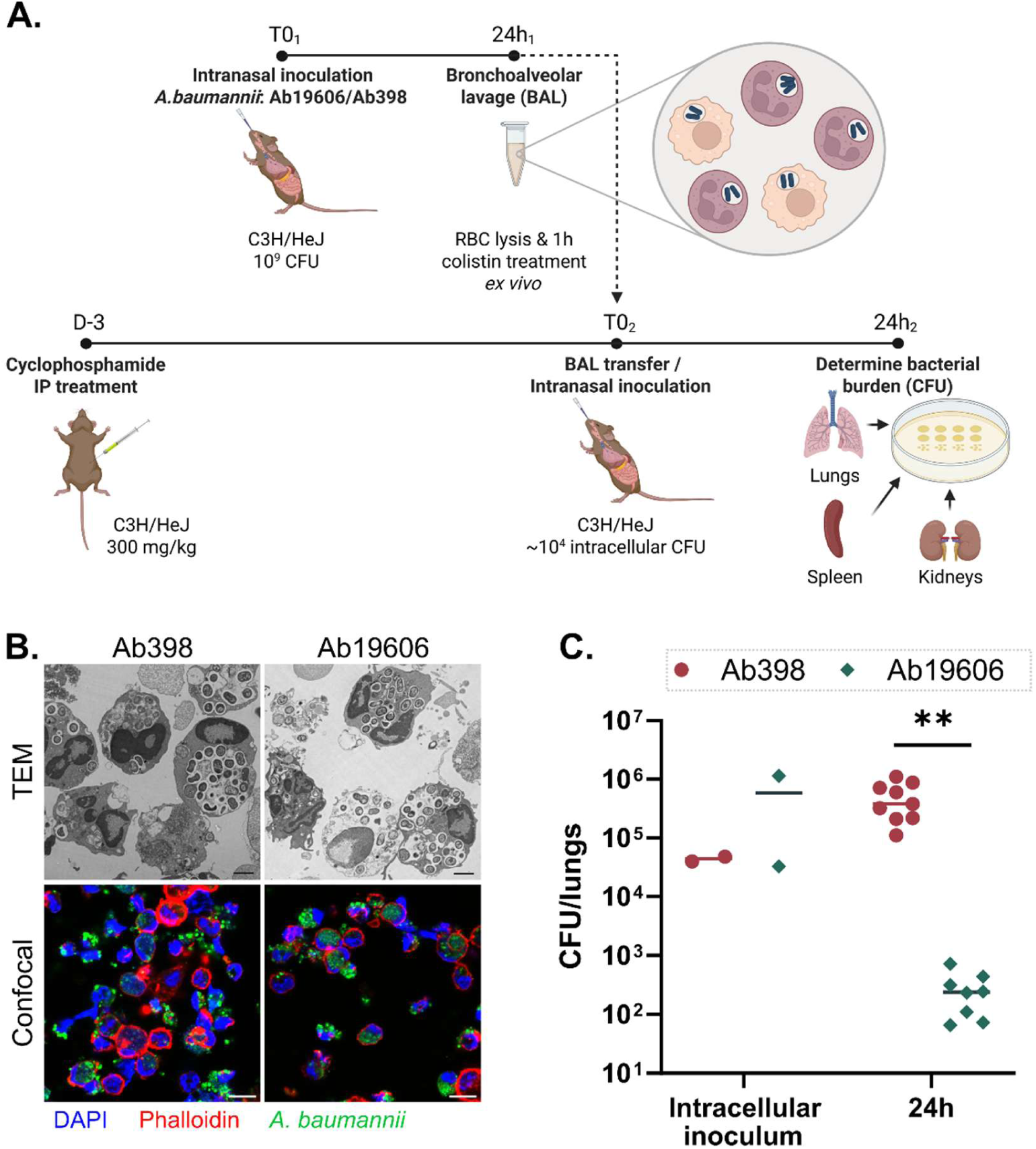
Transfer of BALF cells infected with intracellular *A. baumannii* can seed *de novo* infection in naive cyclophosphamide-treated mice. **A.** Experimental workflow of the BALF cells transfer experiment. RBC: Red blood cells. **B.** Visualization of BALFs following RBC lysis and 1h colistin treatment of C3H/HeJ mice infected with 10^9^ of Ab398 or Ab19606 at 24 hpi by transmission <u>e</u>lectron <u>m</u>icroscopy (TEM – Top) or confocal microscopy (bottom). **C.** Transfer of BAL containing intracellular Ab398 can seed a *de novo* infection in cyclophosphamide-treated C3H/HeJ mice whereas transfer of BAL containing intracellular Ab19606 was readily eliminated from the lungs of mice 24h post infection. Data was statistically analyzed using a t-test with Welch’s correction. **: *p_val_*<0.01.

Colistin-treated BALFs containing approximately 10^4^-10^5^ intracellular bacteria were intranasally transferred to a second group of cyclophosphamide-treated C3H/HeJ mice. Cyclophosphamide, is a potent immunosuppressive agent that induces severe neutropenia, mimicking the condition of highly immunocompromised patients affected by *A. baumannii* infection. It is widely accepted in the field for use in animal models (*33*, *34*). Neutropenia also slows the kinetics of bacterial clearance in the lungs. BALF transferred carried similar amounts of intracellular bacteria for both strains. At 24 hpi, bacterial burdens in lungs, spleens and kidneys of the recipient mice were assessed by CFU enumeration. Mice infected with BALF carrying intracellular Ab398 resulted in a significant increase in bacterial burden in the lung 24 hpi (∼10^5^-10^6^ CFU), indicating that not only intracellular Ab398 are viable, but are also capable of initiating *de novo* lung infection in naive mice (*Figure 1C*). Consistent with the low-dose intranasal inoculation in C3H/HeJ mice, which leads to localized lung infection rather than systemic spread, CFU counts in the spleen and kidneys were undetectable (*17*). Mice that received BALF from Ab19606-infected mice exhibited a significantly lower bacterial burden in their lungs at 24 hpi (∼10^2^). Together, our result demonstrates that intracellular bacteria, at least for some virulent strains such as Ab398, can act as a potentially infectious reservoir.

Upon egress from the host cells, this intracellular bacterial population can seed *de novo* lung infections. Within a single infection, infected BALF cells may serve as a transient niche and reservoir for *A. baumannii* during infection, underscoring the importance of studying its intracellular lifestyle.

### Comparative dual-proteomic analysis

*A. baumannii* lacks the canonical virulence factors (T3SS and T4SS) employed by most intracellular Gram-negative bacteria to survive and replicate within macrophages. To identify factors that promote bacterial survival and replication within cells and contribute to interfere with phagosome maturation, we used a comparative dual-proteomic approach (*Figure 2A*). In this study, we infected mouse J774.A1 macrophages with two replicative *A. baumannii* clinical isolates, Ab398 and Ab803, at a <u>m</u>ultiplicity <u>o</u>f infection (MOI) of 10 (*13*). These strains belong to different <u>m</u>ultilocus <u>s</u>equence typing (MLST) categories and possess different capsule types and LOS outer cores (*Table S1*). Non-infected macrophages and bacteria grown in cell culture media under static conditions served as controls. At 1 hpi, cells were washed and colistin was added to the media for J774.A1-containing samples (infected and uninfected), to kill any extracellular bacteria. At 4 hpi, cells were washed and lysed, and protein content was precipitated. Subsequently, we performed proteomic analyses using a data-independent acquisition (DIA) mass spectrometry approach. Overall, we detected 6,271 and 6,354 macrophage proteins in samples infected by Ab398 and Ab803, respectively (*Figure 2B*). Additionally, we detected 663 and 513 bacterial proteins in samples infected by Ab398 and Ab803, respectively, 452 of which were conserved and detected between the two strains.

**Figure 2.**
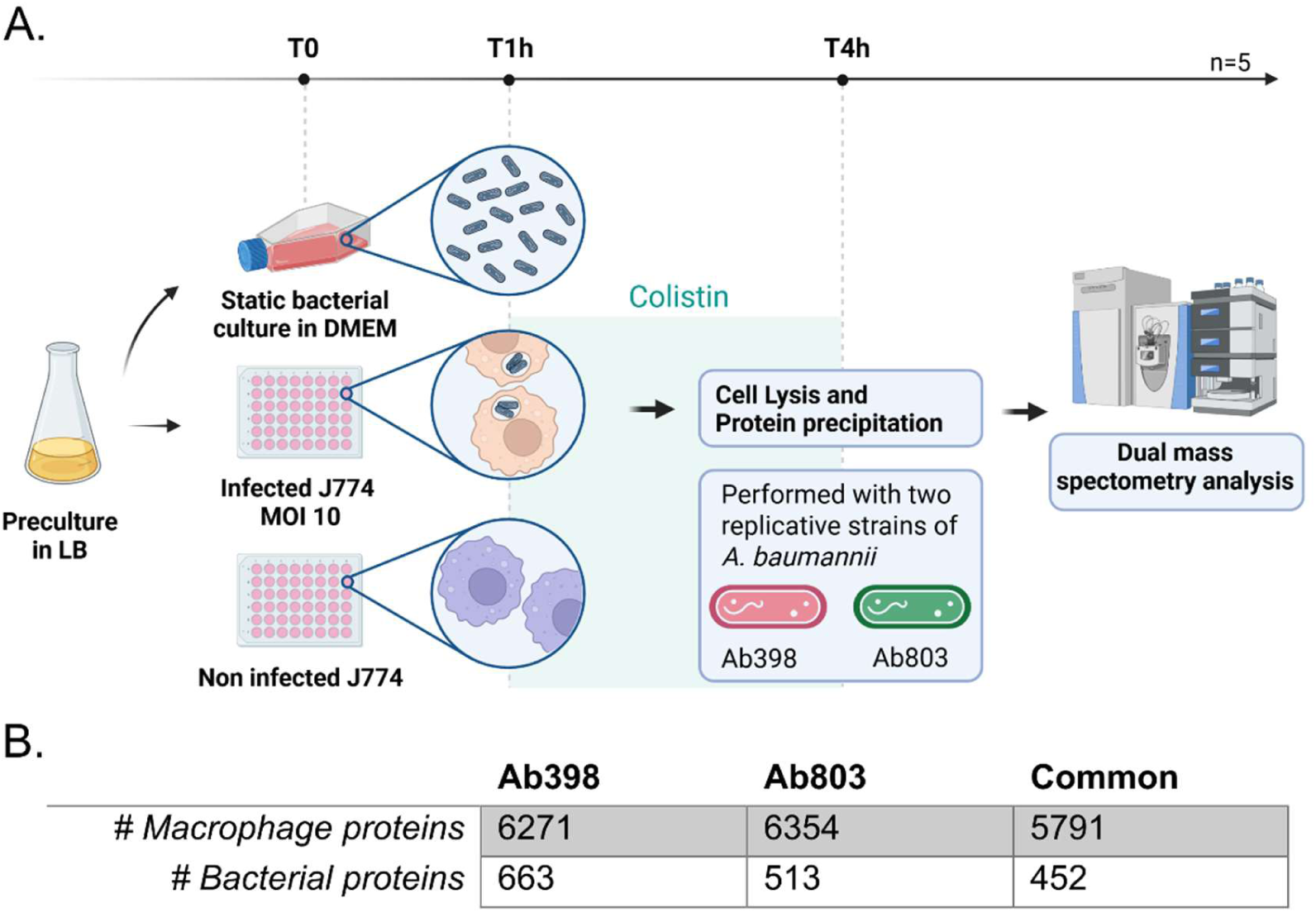
Comparative dual-proteomic analysis of intracellular *A. baumannii* infection of J774.A1 macrophages. **A.** Workflow of the experimental approach. **B.** Number of host and bacterial proteins detected by proteomic analysis across the samples.

### Proteome profiling of *A. baumannii*-infected J774.A1 macrophages

We plotted the Log_2_(Fold Change (FC)) values of host proteomic data following infection by Ab398 against those following infection by Ab803 (*Figure 3A*). The linear distribution of the data along the line of identity indicates a highly similar macrophage response to both *A. baumannii* strains. To explore host proteome remodeling upon intracellular infection, we conducted <u>g</u>ene <u>s</u>et <u>e</u>nrichment <u>a</u>nalysis (GSEA) and calculated <u>n</u>ormalized <u>e</u>nrichment <u>s</u>cores (NES) for hallmark mouse gene sets (*35*). We identified 11-12 enriched gene sets (<u>F</u>alse <u>d</u>iscovery rate (FDR)<0.05), respectively, with the most significant, considering both enrichment score and gene counts being TNFɑ signaling via NF-κB, Inflammatory response, and IFNɣ response (*Figure 3B, Figure S2*). These three pathways share overlapping gene/protein content. Label-free quantitation (LFQ) values for enriched proteins of which levels were significantly altered in at least one strains are presented on *Figure 3C-E,* along with the Log_2_(FC) values for Ab398 and Ab803 infections. Macrophage initially detect lipo<u>o</u>ligo<u>s</u>accharide (LOS) – the major outer membrane component in *A. baumannii* – *via* binding to TLR4, triggering NLRP3 inflammasome activation and IL-1ꞵ production (*36*, *37*). Although NLRP3 levels remained unchanged in our proteomic data, IL-1ꞵ levels significantly increased upon infection (Ab398: Log_2_(FC) = 6.5, *q-val* = 0; Ab803: Log_2_(FC) = 4.2, *q-val* = 4.7E-4). Activated macrophages produce numerous pro-inflammatory cytokine and chemokines including IL-1ꞵ, TNFɑ, Cxcl2, Ccl4, Ccl2 and Cxcl10, all of which were markedly upregulated in infected macrophages in our data (*Figure 3C-E*, *Table S3*, (*10*, *38*)). Additionally, *A. baumannii*-infected macrophages showed increased expression of nitric oxide synthase (NOS2, Ab398: Log_2_(FC)=5.2, *q-val* = 2.4E-4; Ab803: Log_2_(FC)=3.4, *q-val* = 1.3E-3) due to LPS and TNFɑ signaling (*39*, *40*). Similarly, levels of NOD2, an intracellular pathogen recognition receptor sensing *A. baumannii* peptidoglycan, were elevated in Ab398 (Ab398: Log_2_(FC)=1.5, *q-val* = 4.2E-2; (*41*)). Our proteomic findings suggest macrophages undergo pro-inflammatory M1 polarization upon *A. baumannii* infection, however, additional experiments are required to confirm this polarization (*38*). These results are consistent with the findings of Kho et al., 2022 in THP-1 human macrophages infected by *A. baumannii* with or without polymyxin B, suggesting that the pro-inflammatory response observed in our experiments is triggered during initial contact between *A. baumannii* and the macrophages, rather than during its intracellular replication. (*42*).

**Figure 3.**
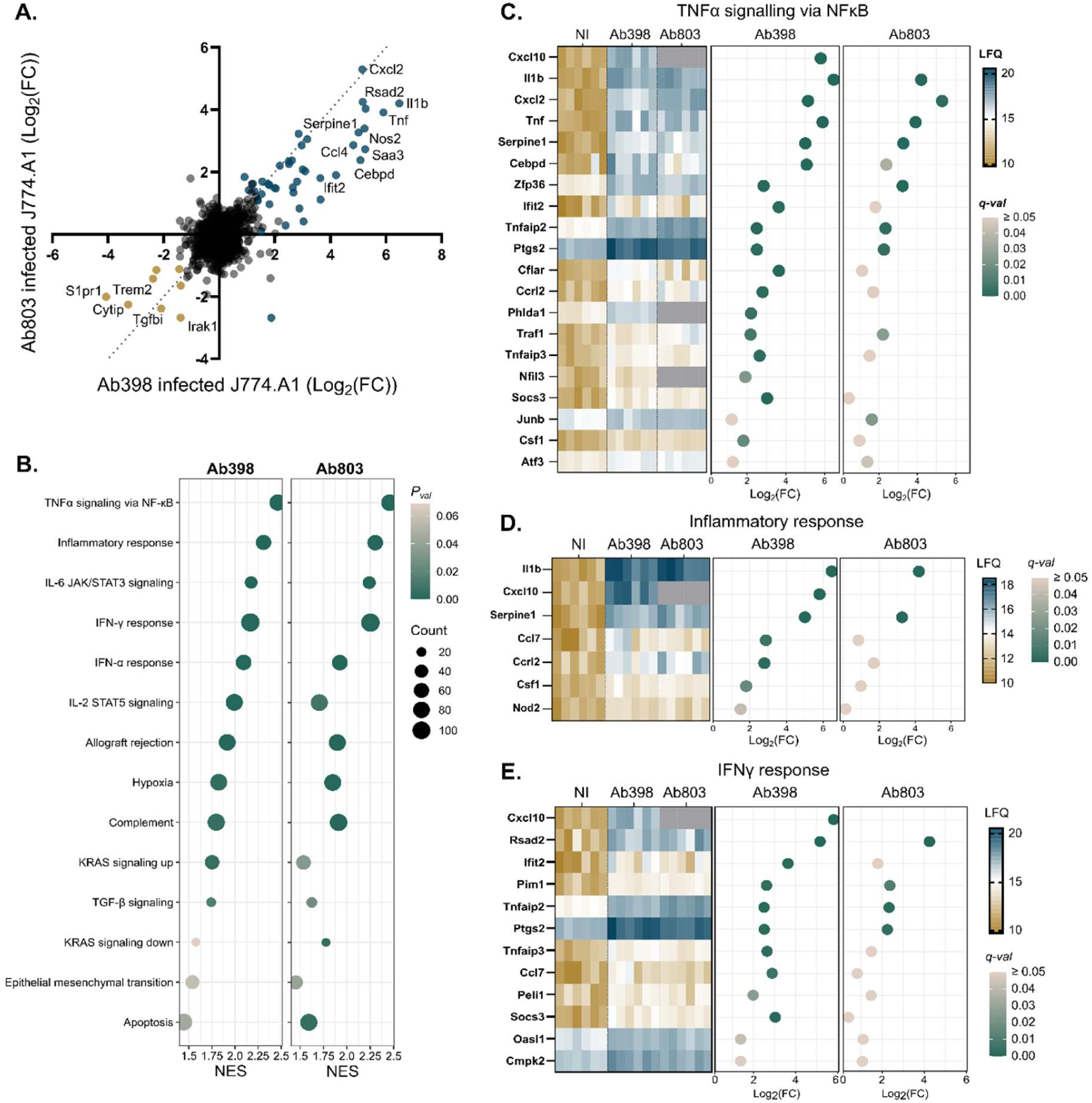
Macrophage proteome profiling upon intracellular *A. baumannii* infection. **A.** Comparison of J774.A1 macrophage response to both *A. baumannii* strains. Blue dots: significantly increased proteins upon infection in at least one of the strains. Yellow dots: significantly decreased proteins upon infection in at least one of the strains. **B.** Gene set enrichment analysis bubble plot. NES: Normalized enrichment score. Count: number of proteins enriched in each gene set. *P_val_*: Nominal p-value. **C-E.** Proteomic normalized label-free quantitation (LFQ) imputed data of main enriched gene sets: TNFɑ signaling via NF-κB (**C.**), Inflammatory response (**D.**) and IFNγ response (**E.**) showing all 6 biological replicates for proteins Log_2_(FC)<0.05 in one or the two strains.

Notably, we observed that the itaconate-synthesizing enzyme Acod1 (IRG1) was significantly induced upon *A. baumannii* infection in J774.A1 macrophages (Ab398: Log_2_(FC)=3.0, *q-val* = 3.7E-4; Ab803: Log_2_(FC)=2.9, *q-val* = 4.8E-4). Itaconate inhibits bacteria species such as *Mycobacterium tuberculosis* and *Salmonella,* while others like *Pseudomonas aeruginosa* and *Staphylococcus aureus* can exploit it as a source of nutrients (*43–47*). To our knowledge, Acod1 induction has not been previously reported for infection by *A. baumannii*. We confirmed this finding by immunoblot (*Figure S3*). In our analysis we included the non-replicative strain Ab19606, which also induced Acod1/IRG1, suggesting that induction of Acod1 is not a response specific to intracellularly replicating *A. baumannii*, consistently with its induction via the TLR4/NF-кB signaling pathway (*48*). To test whether intracellular Ab19606 could be eliminated due to a higher sensitivity to itaconate, we added itaconic acid (IA) in culture media. Although neither Ab19606, Ab398 nor Ab803 strains were able to use IA as a sole carbon source (*Figure S4A*), none of the strains’ growth was affected by the addition of IA to the media (*Figure S4B-D*).

Taken together, intracellular *A. baumannii* infection triggers a macrophage inflammatory response, characterized by the production of pro-inflammatory cytokines and chemokines, and the induction of Acod1.

### Proteomic profiling of intracellular *A. baumannii*

Two intracellular *A. baumannii* strains, Ab398 and Ab803 were used in our dual comparative proteomic analysis of infected J774.A1 macrophages. Both strains survive and replicate within ACVs in J774.A1 macrophages (*13*). Proteomic analysis was conducted at 4 hpi as the peak of replication typically occurs between 4 and 8 hpi in J774.A1 cells. Bacterial cultures grown under static condition for 4h in DMEM + 10% <u>h</u>eat inactivated <u>F</u>etal <u>B</u>ovine <u>S</u>erum (hi-FBS) - the same media used for infection – served as baseline. Input samples grown in LB were also analyzed but excluded from the final analysis, as the bacterial growth state following a 16-hour pre-culture differs significantly from the tested conditions. To allow comparison between sample groups *A. baumannii* proteins observed during infection were normalized to the median of the LB groups as shown in *Figure S5*. *A. baumannii* exhibits extensive genetic diversity, with a large pan-genome exceeding 20,000 genes. As a result, not all genes are conserved between the two strains. *Figure 4A* shows the Log_2_(fold change (FC)) of all 452 conserved proteins detected in both strains. Intracellular infection of J774.A1 macrophages induced highly similar changes in *A. baumannii* protein levels across these conserved proteins, as most data aligned closely with the line of identity. Among these 452 bacterial proteins, 73 exhibited significant alterations in abundance during intracellular infection compared to their respective control (static growth in culture media).

**Figure 4.**
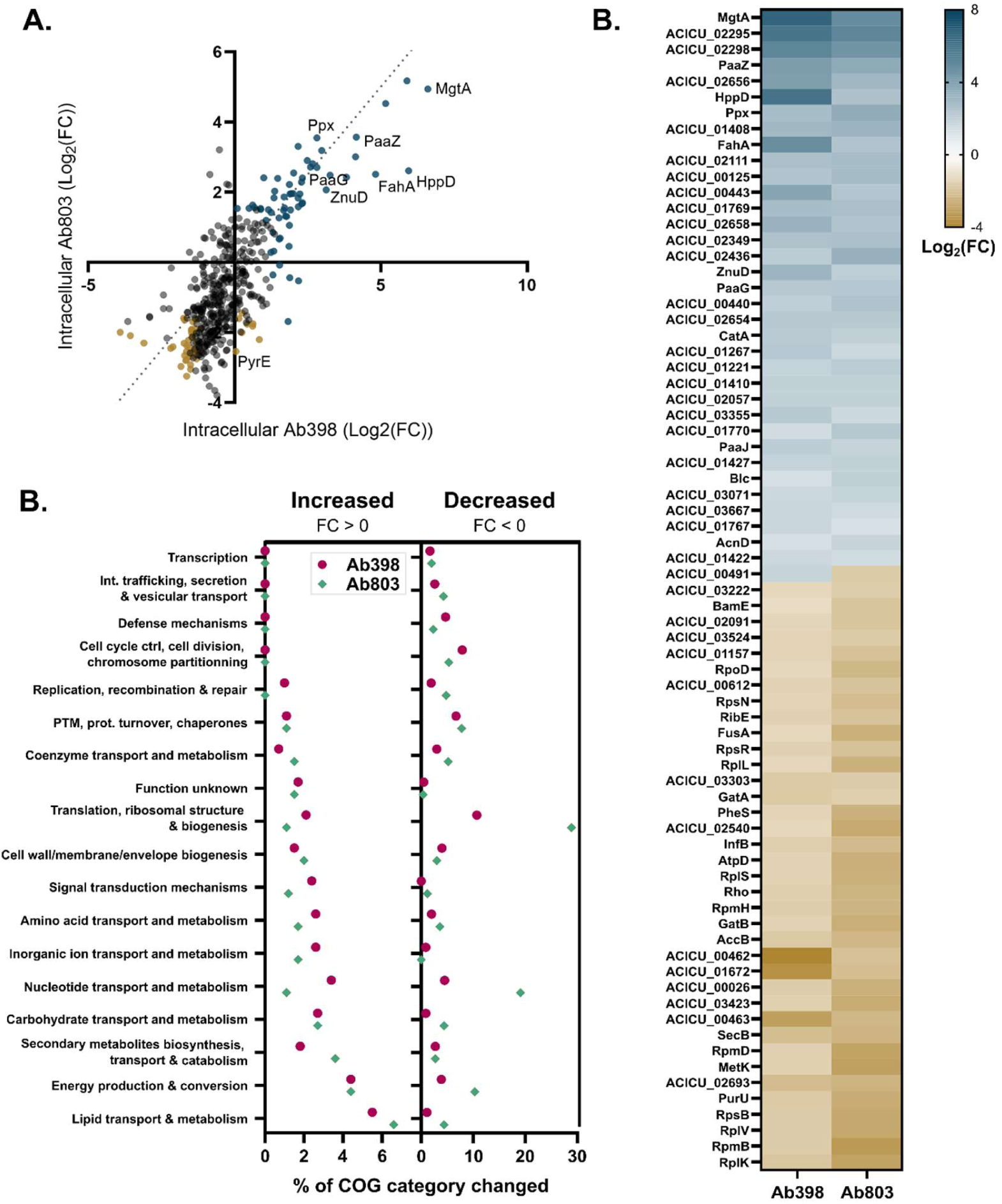
Proteome profiling of intra-macrophage *A. baumannii*. **A.** Comparison of *A. baumannii* proteome change during intracellular J774.A1 macrophage infection in Ab398 *vs* Ab803. Blue dots: significantly increased proteins upon intracellular infection in at least one of the strains. Yellow dots: significantly decreased proteins upon intracellular infection in at least one of the strains. Only conserved proteins which were detected in the proteomic analysis in both strains are plotted. **B.** COG pathway analysis. The percentage of proteins significantly changed within a COG pathway; either increased (Fold change (FC) > 0) or decreased (FC < 0) in each strain. **C.** Heat map showing significantly changed proteins in the two strains following J774.A1 macrophage infection based on their Log_2_(FC).

Significantly altered proteins belonged to diverse pathways, as demonstrated by the analysis of clusters of orthologous groups (COG) categories (*Figure 4B*). In both strains, a higher proportion of proteins involved in ‘lipid transport and metabolism’ and ‘energy production and conversion’ increased during macrophage infection. Conversely, proteins that decreased during infection were primarily associated with ‘translation, ribosomal structure and biogenesis’, ‘nucleotide transport and metabolism’ or even ‘cell cycle control, cell division, chromosome partitioning’. *Figure 4C* shows the most significantly altered proteins on both strains.

### Analysis of intra-macrophage *A. baumannii proteome*

The proteomic results for the bacterial proteins are presented in *Table S4*. Several proteins involved in defense against host nutritional immunity were increased in intracellular bacteria. Hosts imposed nutritional immunity limits bacterial access to trace metals, such as Zinc (Zn^2+^), Magnesium (Mg^2+^), and Iron (Fe^2+^), preventing their proliferation during infection (*49*). Bacteria counteract by inducing the corresponding transporters. Our proteomic data showed increased levels of both ZnuD and MgtA following macrophage infection (*Table 1*). In *A. baumannii*, Zn^2+^ is imported by the Zn^2+^ uptake (ZnuABCD) system, which contributes to bacterial survival in *in vivo* infections (*50–52*). Similarly, MgtA type Mg^2+^ transporters, extensively studied in *Salmonella*, play a crucial role in virulence in mice and replication in macrophages (*53–55*).

**Table 1:** Selected altered *A. baumannii* proteins following intracellular macrophage infection. Deletion mutants in genes corresponding to the proteins in bold were used for further analysis. NS: non-significant. ND: non detected.

| Protein name | Ab398 |  | Ab803 |  |  |
| --- | --- | --- | --- | --- | --- |
|  | Log<sub>2</sub>(FC) | q-value | Log<sub>2</sub>(FC) | q-value |  |
| **MgtA** | 6.60 | 0 | 4.94 | 1.33E-03 | Nutritional immunity |
| **ZnuD** | 3.13 | 0 | 2.06 | 1.64E-03 |  |
| PaaZ | 4.15 | 0 | 3.57 | 1.70E-03 | Stress response |
| **PaaG** | 2.32 | 4.44E-04 | 2.40 | 8.42E-04 |  |
| PaaJ | 2.17 | 3.33E-03 | 1.83 | 7.10E-03 |  |
| **Ppx** | 2.81 | 1.81E-03 | 3.55 | 2.15E-03 |  |
| Ppk1 | -1.31 | 6.22E-02 | -2.02 | 1.86E-02 |  |
| PstS | 2.29 | 0 | 2.29 | 7.44E-04 |  |
| SodB | 1.86 | 3.47E-03 | NS |  |  |
| KatG | 1.54 | 3.45E-02 | NS |  |  |
| **LpsB** | 2.02 | 4.12E-03 | 2.55 | 1.76E-02 | Surface modification |
| Blc | 1.37 | 4.04E-02 | 2.03 | 3.95E-03 |  |
| **FahA** | 4.82 | 0 | 2.51 | 8.00E-04 | Energy source utilization |
| **Ald1** | 5.16 | 0 | 4.53 | 0 |  |
| CatA | 2.23 | 2.90E-03 | 1.95 | 4.19E-03 |  |
| NuoE | 2.86 | 0 | ND |  |  |
| MmsB | 2.09 | 3.53E-03 | ND |  |  |

Proteins encoded by the <u>p</u>henyl<u>a</u>cetic <u>a</u>cid (PAA) catabolism operon were also increased upon intracellular infection, including PaaZ, PaaG and PaaJ (*Table 1*). In *A. baumannii*, this pathway is involved in stress-response and virulence. PAA is an intermediate in the catabolism of phenylalanine. In *A. baumannii*, the virulence of a mutant unable to catabolize PAA was significantly impaired in both murine and zebrafish models of infection (*56*, *57*). Our analysis also revealed increased levels of exopolyphosphatase (Ppx), responsible for the catabolism of polyphosphate (polyP), while levels of Ppk1, the polyphosphate kinase synthetizing polyP, were decreased upon infection. PolyPs are linear chains of phosphate residues linked by high-energy bonds, which can be stored in polyP granules and play an important role in the virulence and stress response in many bacterial species, including *P. aeruginosa* and *A. baumannii* (*58–61*). Further highlighting the requirements for phosphate in *A. baumannii* intracellular survival/replication, protein levels of PstS, a phosphate transporter, were also increased in both strains (*62*). Other stress responses were activated such as response to the oxidative stress through increase of the superoxide dismutase (SodB) and the catalase (KatG) in the Ab398 strain.

In addition, proteins levels of LpsB, a glycosyl transferase required for the addition of outer core to the nascent lipo<u>o</u>ligo<u>s</u>accharide (LOS), were increased (*63*). This is consistent with previously reported reduced virulence of *lps*B mutants in serum and *in vivo* rat and mouse models (*64–66*). Also involved in bacterial cell surface modification during the intracellular lifestyle, our proteomic analysis showed increased levels of bacterial lipocalin Blc in both strains, an inner membrane protein expressed in response to starvation or high osmolarity, allowing maintenance of membrane and envelope integrity (*67*, *68*).

Multiple proteins involved in metabolism and energy source utilization were also increased upon macrophage infection, including FahA and Ald1. FahA is a fumarylacetoacetase (also known as HmgC), belonging to the homogentisate (Hmg) degradation pathway (*69*, *70*). Together with the PAA catabolism pathway, the Hmg pathway participates in the degradation of the amino acids phenylalanine and tyrosine. Ald1 is annotated as a long-chain aldehyde dehydrogenase, which could play a role in the catabolism of n-alkanes or ethanolamine utilization (*71–73*). In line with these findings, in both *Listeria monocytogenes* and *Salmonella enterica* serovar Typhimurium, intracellular replication of mutants in the <u>e</u>thanolamine <u>ut</u>ilization (eut) pathway was impaired (*74–76*). *A. baumannii* also appears to rely at least partly on intracellular ethanolamine as a source of carbon and nitrogen. Other proteins induced were CatA, a protein involved in aromatic compound metabolism (*77*), the NADH-quinone oxidoreductase subunit E (NuoE), and the hydroxyisobutyrate dehydrogenase MmsB in Ab398.

Overall, the analysis of the proteins with known function increased in intracellular bacteria were mainly involved in four main functional categories: counteracting nutritional immunity, stress responses, surface modification, and energy source utilization (*Table 1*). Our analysis additionally identified many proteins of unknown function that are up- or down-regulated upon macrophage infection (*Table S4*). The role of these proteins will be the scope of future work.

### *A. baumannii* survival and replication in macrophages is multifactorial

We engineered Ab398 deletion mutants in genes corresponding to the most altered proteins in each of the identified functional categories. No significant growth defects were observed when these mutants were grown in DMEM + hi-FBS (*Figure S6A*). We subsequently assessed the impact of these mutations on replication within J774.A1 macrophages over 24h, respective to that of the wild type Ab398 strain (*Figure 5A-H*). The complete summary of the data is shown in *Figure S7*. To complement the analysis, we conducted confocal microscopy analysis of J774.A1 infected with each mutant at 24 hpi (*Figure 5I*). All the mutants displayed impaired replication ability in macrophages to different extents. Both mutant strains involved in nutritional immunity (Δ*mgtA* and Δ*znuD*) displayed moderate but statistically significant and reproducible replication defects at all times, with the Δ*mgtA* showing a more dramatic effect at 12 hpi. This aligns with findings in *Salmonella*, in which *mgtA* and *mgtCB* are required a later time during macrophage infection (*54*). Δ*paaG* and Δ*ppx* mutants showed markedly reduced replication while Δ*fahA* and Δ*ald1* mutants displayed little to no replication in J774.A1 macrophages. Interestingly, an *lpsB-*deficient mutant, in which we could confirm the absence of LOS outer core (*Figure S8*), was unable to survive in macrophages and was readily eliminated over time. We showed that this was not due to increased sensitivity to acidic pH as the Δ*lpsB* mutants grow similarly to the wildtype strain in LB buffered at pH5 (*Figure S6B*). In line with data obtained by antibiotic protection assays, confocal microscopy showed a significant reduction in intra-macrophage bacterial burdens (*Figure 5C* – green signal), with the most affected mutants being Δ*lpsB* and Δ*ald1*.

**Figure 5.**
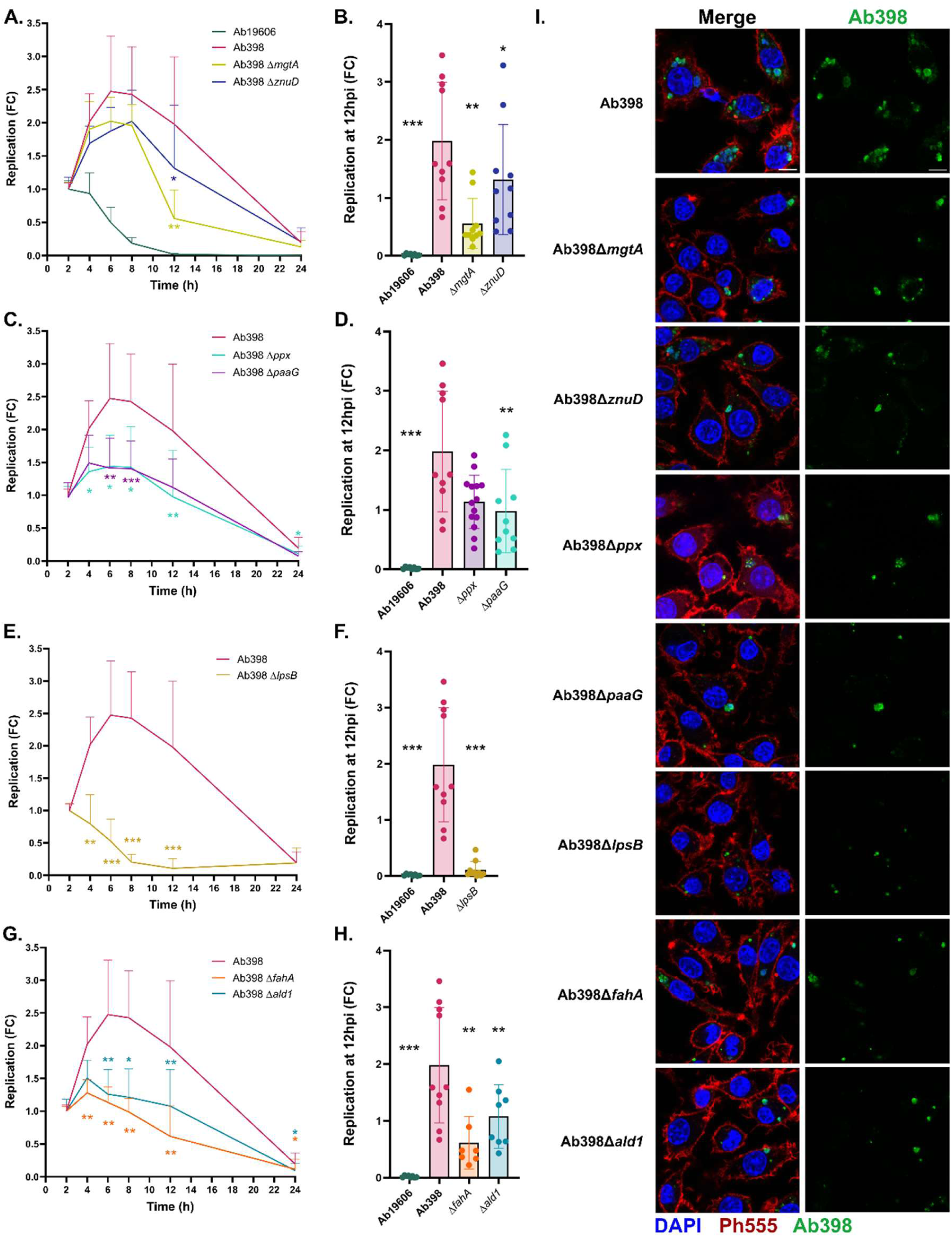
*A. baumannii* intracellular survival and replication relies on multiple strategies. **A, C, E, G.** Bacterial replication within J774.A1 macrophages. Antibiotic protection assay performed with an MOI of 10 over 24h. **B, D, F, H.** Individual experimental data at 12 hours post infection (hpi) with Ab398, Ab19606, used as non-replicative control and Ab398Δ*mgtA and* Ab398Δ*znuD* (**A-B**), Ab398Δ*ppx* and Ab398Δ*paaG* (**C-D**), Ab398Δ*lpsB* (**E-F**), Ab398Δ*fahA* and Ab398Δ*ald1* (**G-H**). FC: fold change compared to 2 hpi. One-way ANOVA was performed on log-transformed data and compared to Ab398. *: *P_val_* < 0.05, **: *P_val_* < 0.01, ***: *P_val_* < 0.001. **I.** Confocal microscopy of Ab398 and deletion mutants at 24 hpi. Macrophages were infected at an MOI of 10 and fixed in 4% PFA at 24 hpi. Samples were permeabilized and bacteria were immunolabeled with anti-*A. baumannii* primary antibodies and anti-rabbit Alexa 488 secondary antibody (Green), and J774.A1 cells nuclei were labeled with DAPI (Blue), and actin was labeled with Phalloidin 555 (Red). Merged color channels are shown on the left and the panels on the right show only Ab398.

Collectively, these results reveal that *A. baumannii* employs a multifactorial strategy to survive and replicate within macrophages. This strategy includes counteracting host-imposed nutritional immunity and activating mechanisms of stress response (including polyP and PAA catabolism) and adapting to its intracellular niche through the utilization of ethanolamine and increased catabolism of tyrosine and phenylalanine. Furthermore, our experiments show that an intact LOS is required for intracellular survival within macrophages.

## Discussion

*A. baumannii* is a highly successful nosocomial pathogen, with an alarming increase in antibiotic resistance rates, and a high mortality among the immunocompromised patients it often infects. Understanding *A. baumannii* pathogenesis is therefore crucial for adapting treatment options. In this study, by employing BALF transfer following *ex vivo* antibiotic treatment, we found that intracellular *A. baumannii* is capable of seeding *de novo* lung infection in naive mice, highlighting the biological significance of *A. baumannii’s* intracellular lifestyle. Comparative dual-proteomic analysis of J774.A1 macrophages infected with replicative strains of *A. baumannii* enabled the molecular characterization of the *A. baumannii*-macrophage interaction interface. *Figure 6* provides a summary of our key findings. Macrophages activated classical antibacterial pathways, including the inflammatory response, the TNFα and Type I interferon signaling pathways. We demonstrate that *A. baumannii* infection leads to the production of Acod1/IRG1. Moreover, we show that *A. baumannii* uses a multifactorial strategy to survive and replicate within macrophages, relying primarily on repurposed molecular determinants and stress response mechanisms that enable it to thrive within the ACV.

**Figure 6:**
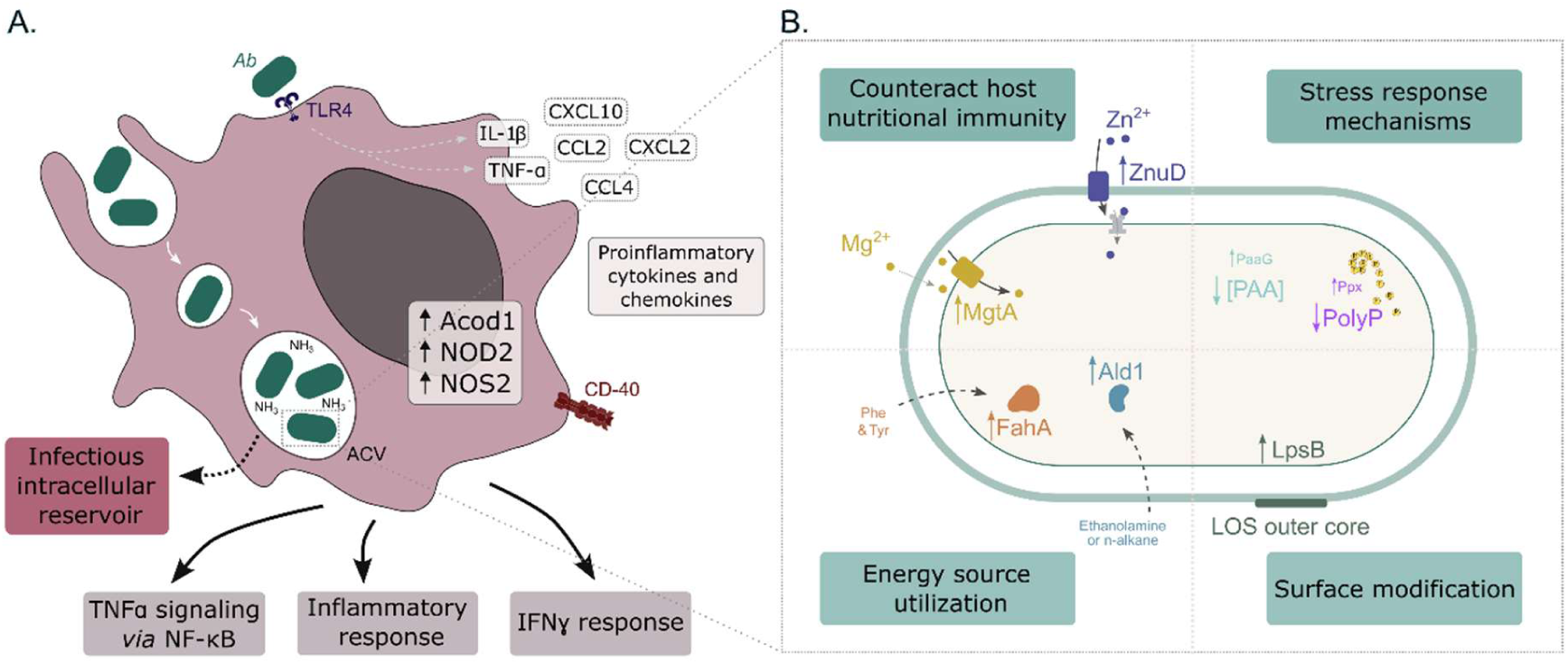
Proposed model of the molecular interface between intracellular *A. baumannii* and macrophages. **A**. Replicative *A*. *baumannii* is phagocytosed by macrophages and establishes the *Acinetobacter*-containing vacuole (ACV). Bacterial replication produces ammonia which neutralizes the luminal microenvironment of the ACV. Intracellular infection of macrophage triggers the activation of TNFα signaling via NF-кB, which leads to an inflammatory response and IFNγ response, releasing proinflammatory cytokines and chemokines. *In vivo*, intracellular bacteria were shown to act as reservoir for lung infection. **B**. *A. baumannii* thrives intracellularly employing a ‘persist and resist’ strategy such as counteracting host imposed nutritional immunity, stress response mechanisms, shifting energy source utilization and modifying its surface.

TNFɑ signaling *via* NF-κB is a key component of the bacterial-induced immune response, and we show that this is also the case for *A. baumannii* infection. Our proteomic analysis was conducted on two replicative *A. baumannii* strains. One limitation of our study is that we cannot fully discriminate between the host response triggered by initial bacterial contact and the response to intracellular replication. Furthermore, the impact of host immune activation induced by *A. baumannii* on its intracellular phase remains unclear. Some pathogens exploit host immune responses to their advantage. For instance, in *P. aeruginosa,* activation of the NF-кB signaling pathway in host A549 cells supports intracellular bacterial survival (*78*). Further studies, including with non-replicative *A. baumannii strains,* and analyses at different timepoints, will be required to dissect the specific contribution of each step to the macrophage’s immune response and the impact of the host response on *A. baumannii* replication.

Our study did not reveal obvious virulence factors or specialized toxins for intracellular replication, unlike other pathogens such as Salmonella, which utilize T3SS effectors (*79*). Instead, we identified upregulation of proteins involved in multiple adaptive mechanisms, including overcoming nutritional immunity (MgtA and ZnuD), known stress response mechanisms (PAA catabolism and polyP), modification of the bacterial surface (LpsB), and metabolic switching/energy source utilization (FahA and Ald1). Additionally, previous studies have shown that *A. baumannii* produces high levels of ammonia, a byproduct of amino acid catabolism, which plays a critical role in neutralizing acidic pH within ACVs (*13*). Collectively, these findings suggest that *A. baumannii* employs a ‘persist and resist’ strategy to survive and replicate within macrophages, mirroring its adaptation to other hostile environments (*80*). Multiple proteins of unknown function were identified to be induced or repressed (*Table S4*). Some of these may contribute to *A. baumannii’s* intracellular replication, which will be the focus of future work.

In response to bacterial infection, host-imposed nutritional immunity typically depletes iron from the phagolysosomal compartment in a similar manner. We demonstrated that *A. baumannii* combats the nutritional immunity imposed by the host by upregulating the production of trace metal transporters (MgtA and ZnuD). Surprisingly, we did not detect proteins involved in iron scavenging. In fact, levels of several siderophore-associated proteins, usually produced and secreted in iron-limiting conditions, were decreased in Ab398, during macrophage infection (*Table S2*). A plausible hypothesis is that *A. baumannii* may be altering their selection of siderophores (*81*, *82*). However, our proteomic analysis did not detect increased proteins associated with siderophore production. Alternatively, the rapid pH alkalization of the ACV caused by the ammonia secreted by bacteria may somehow lead to increased iron availability in the ACV. Investigating this mechanism will be a key focus of future research. *A. baumannii* can establish long-term infections, with an average hospital stay of 30 days for patients with lung infections, a duration that increases in cases involving multidrug-resistant strains (*83–85*). Complicating treatment further, *A. baumannii* frequently causes recurrent and recalcitrant infections, which can be partially attributed to the bacterium’s remarkable ability to withstand antibiotic treatment and survive under various stress conditions (*86–89*). Our data suggest that macrophages serve as a transient niche and reservoir for *A. baumannii* during infection (*Figure 1,* (*17*)). It is tempting to speculate that through repeated cycles of invasion, intracellular replication, and lytic egress from macrophages and neutrophils, *A. baumannii* gains additional protection from both the impaired immune defenses of immunocompromised hosts and antibiotics that poorly penetrate host cell membranes, thereby enhancing its infective capacity and treatment failure (*19*, *90*)). We previously demonstrated that intracellular *A. baumannii* residing in the bladder epithelium of mice after a UTI can act as a reservoir for resurgent infections (*23*). Whether *A. baumannii* similarly establishes reservoirs in lung epithelial cells remains to be determined. Our findings support the idea that the intracellular lifestyle of *A. baumannii* may contribute to recalcitrant infections and should be considered when developing new treatment strategies.

## Material and Methods

### Bacterial strains, culture conditions

Plasmids and strains used in this work are listed in *Table S5*. Bacterial cultures were grown at 37°C in Lennox broth/agar (LB) under agitation. Cultures and plates were supplemented with 10 μg/mL chloramphenicol, 50 μg/mL apramycin, or 10% sucrose when appropriate.

### Bacterial genetic manipulations

Deletion mutants were engineered by amplifying approximately 500-1000 bp upstream and downstream flanking regions of the gene of interest from genomic DNA by PCR using appropriate primer pairs (F1/R1, F2/R2). Independently, pEX18-Apr was amplified using F-pEX18-cloning and R-pEX18-cloning primers. All plasmids and primers used in this work are listed in *Tables S5-6*. Subsequently, overlapping fragments were cloned into linearized pEX18-Apr using NEBuilder HiFi DNA Assembly Master Mix (NEB). Plasmid were electroporated into *E. coli* and plasmid sequence was verified by sequencing (Plasmidsaurus). Using quadruparental mating, allelic exchange vectors were introduced into *A. baumannii*, with pRK2013 and pTNS2 as a helper plasmid, as previously described (*91*, *92*). Selection of merodiploid clones resulting from homologous recombination was performed on LB agar plates containing chloramphenicol and apramycin. Counterselection for double crossover was performed by streaking these colonies on LB agar without NaCl supplemented with 10% sucrose and grown at room temperature for 48h. Mutants were then confirmed by PCR analyses and whole-genome sequencing. GFP-expressing strains were generated by the integration of the vector pUC18TminiTn7T-Apr::gfpd2 using a quadruparental conjugation technique. Selection was achieved using LB supplemented with apramycin and chloramphenicol, and insertion at the mTn7 site was confirmed by PCR analyses.

### Cell culture conditions

The J774A.1 mouse macrophage cell line (ATCC TIB-67) was cultured in Dulbecco’s Modified Eagle Medium (DMEM) High Glucose (Sigma-Aldrich) supplemented with 10% heat-inactivated Fetal Bovine Serum (hi-FBS, Corning) at 37°C and 5% CO2.

### Murine model of bronchoalveolar lavage transfer

All animal experiments were approved by the Washington University Animal Care and Use Committee, and we have complied with all relevant ethical regulations. For BALF transfer, 3 days before infection, 6- to 8-week-old female C3H/HeJ (Jackson Laboratory, Bar Harbor, ME) mice were treated intraperitoneally with 100 μL of 300 mg/kg (6 mg/mouse) of cyclophosphamide (TCI), dissolved in PBS and filtered. On the day of the first infection, overnight bacterial cultures were sub-cultured at a 1:200 dilution in fresh LB and grown with shaking at 37°C for 3 h to mid-exponential growth phase. Cultures were washed twice with PBS and normalized to OD_600nm_=100. Naive C3H/HeJ mice were anesthetized with 4% isoflurane and were intranasally inoculated with 10^9^ CFU (35µl of culture). At 24 hpi, mice were sacrificed by isoflurane overdose and <u>b</u>roncho<u>a</u>lveolar lavage (BAL) was collected in 1 mL of PBS-EDTA 1 mM. <u>B</u>AL fluids (BALFs) were pooled followed by red blood cell lysis by incubation for 5 min on ice in 5 mL of Pharm Lyse Buffer diluted in PBS (BD Biosciences 555899), and the reaction was stopped by the addition of 30 mL of PBS. Cells were then incubated for 1h in DMEM 20% hi-FBS with colistin 50 μg/mL for Ab398 and 100 μg/mL for Ab19606 (Minimal inhibitory concentration (MIC) twice higher than that of Ab398) under agitation to eliminate all extracellular bacteria. Cellular content was centrifuged at 600 xg for 6 min, washed twice with PBS and resuspended in PBS (10 BALFs pooled, treated and resuspended in 300 μL). Subsequently, cyclophosphamide-treated mice were intranasally inoculated with 35 μL of the pooled treated BALF. Intracellular bacterial count was determined by lysis of the host cells by Triton X-100 (0.025%) and serial dilutions of bacterial suspensions were plated for CFU counting. At 24 hpi, mice were sacrificed, and CFU in the lungs, spleen, and kidneys were quantified by serial dilution plating the homogenized organs. Additionally, half of the lung slurry was plated onto LB chloramphenicol plate to decrease the bacterial detection limit.

### Transmission electron microscopy (TEM) of antibiotic-treated BALF

C3H/HeJ female mice were intranasally inoculated with 10^9^ CFUs of either Ab398 or Ab19606 strain. At 24 hpi, mice were sacrificed by isoflurane overdose and BAL was performed with 1 mL of PBS-EDTA 1 mM. For ultrastructural analyses, colistin-treated BALF (100 μg /mL) samples were gently pelleted and fixed in 2% paraformaldehyde/2.5% glutaraldehyde (Ted Pella Inc., Redding, CA) in 100 mM cacodylate buffer, pH 7.2 for 2 h at room temperature. Samples were washed in cacodylate buffer and postfixed in 1% osmium tetroxide (Ted Pella Inc.)/ 1.5% potassium ferricyanide (Sigma, St. Louis, MO) for 1h. Samples were then rinsed extensively in dH_2_0 prior to ‘en bloc’ staining with 1% aqueous uranyl acetate (Ted Pella Inc.) for 1h. Following several rinses in dH_2_0, samples were dehydrated in a graded series of ethanol and embedded in Eponate 12 resin (Ted Pella Inc.). Ultrathin serial sections of 95 nm were cut with a Leica Ultracut UCT ultramicrotome (Leica Microsystems Inc., Bannockburn, IL), stained with uranyl acetate and lead citrate, and viewed on a JEOL 1200 EX transmission electron microscope (JEOL USA Inc., Peabody, MA) equipped with an AMT 8-megapixel digital camera and AMT Image Capture Engine V602 software (Advanced Microscopy Techniques, Woburn, MA).

### Antibiotic protection assay

Bacterial intracellular replication was performed as previously described (*13*). Briefly, J774A.1 cells were seeded 16 h before the experiment in 48-well plates (3x10^5^ cells/well). *A*. *baumannii* overnight cultures were inoculated from fresh LB-chloramphenicol plates. Bacterial cultures were pelleted for 5 min at 6500 rpm and washed twice with PBS. Bacterial cultures were normalized to OD_600nm_ = 1 and appropriate volumes were added to 500 μL of DMEM high glucose (Sigma-Aldrich), 10% hi-FBS per well to achieve an MOI = 10. Infected cells were centrifuged for 10 min at 200 xg and incubated at 37°C and 5% CO2. 1 hpi, macrophages were washed thrice with PBS and treated with DMEM supplemented with 10% FBS and 50 μg/mL of colistin to eliminate extracellular bacteria. At the indicated time points, cells were washed thrice with PBS and lysed with 500 μL of Triton X-100 (0.025%) per well. Serial dilutions of bacterial suspensions obtained were plated. Bacterial replication was assessed by CFU counting. Replication fold change (FC) for each timepoint was calculated by ratio compared to the average of the two technical duplicates at the 2-h timepoint. Data was collected from at least 3 independent experiments.

### Dual comparative proteomic analysis

#### A. baumannii infection

Antibiotic protection assays were performed as described above using a full 48-well plate per conditions. At 4 hpi, plates were washed 3 times with PBS and 100 μL of Triton X-100 (0.025%) was added per well. After a few minutes, samples were collected in 5X lysis buffer (20% SDS, 500mM Tris pH 8.5). All lysed samples were boiled for 10min. Protein concentration was measured for each sample with the DC protein assay kit (Bio-Rad) and visualized by SDS-PAGE, prior to addition of 10 mM of dithiothreitol (DTT) to all the samples.

#### Acetone precipitation

1 mg of protein of each sample was used for acetone precipitation. Following the addition of 100 mM of NaCl to the sample, 4 volumes of ice-cold acetone were added, and samples were placed at -20°C overnight. Samples were then centrifuged at 4000 xg for 10 min at 4°C. The pellets were resuspended by vortexing in Milli-Q water and 4 volumes of ice-cold acetone were added. Samples were incubated at -20°C for at least 4h. Precipitated proteins were harvested by centrifugation at 6000 xg for 15 min at 4°C and pellets were air dried.

#### Sample preparation for Proteomic analysis

Acetone precipitated samples were solubilized in 4% SDS, 100 mM Tris pH 8.5 by boiling for 10 min at 95°C, and then protein concentrations were quantified using bicinchoninic acid assays (Thermo Fisher Scientific). 200 μg of protein for each replicate was prepared for protein digestion using S-trap mini columns (Protifi, USA) according to the manufacturer’s instructions. Briefly, samples were reduced with 10 mM DTT for 10 mins at 95°C and then alkylated with 40 mM Iodoacetamide in the dark for 1 h. Reduced and alkylated samples were then acidified with phosphoric acid to a final concentration of 1.2% then diluted with seven volumes of S-trap wash buffer (90% methanol, 100 mM Tetraethylammonium bromide pH 7.1). Samples were loaded onto S-trap mini columns and washed 4 times with S-trap wash buffer. Samples were then digested with 4μg of Trypsin/Lys-C (Promega) overnight before being collected by centrifugation with washes of 100mM Tetraethylammonium bromide, followed by 0.2% formic acid followed by 0.2% formic acid / 50% acetonitrile. Samples were dried down and further cleaned up using C18 Stage (*93*, *94*) tips to ensure the removal of any particulate matter.

#### Reverse phase Liquid chromatography–mass spectrometry

Prepared purified peptides from each sample were resuspended in Buffer A* (2% acetonitrile, 0.01% trifluoroacetic acid) and separated using a two-column chromatography setup composed of a PepMap100 C_18_ 20-mm by 75-mm trap and a PepMap C_18_ 500-mm by 75-mm analytical column (Thermo Fisher Scientific). Samples were concentrated onto the trap column at 5 ml/min for 7 min with Buffer A (0.1% formic acid, 2% DMSO) and then infused into an Orbitrap Exploris 480 (Thermo Fisher Scientific) at 300 nl/minute via the analytical column using a Dionex Ultimate 3000 UPLC (Thermo Fisher Scientific). Samples were separated using a 95-minute analytical gradient undertaken by altering the buffer composition from 4% Buffer B (0.1% formic acid, 77.9% acetonitrile, 2% DMSO) to 25% B over 75 min, then from 25% B to 40% B over 4 min, then from 40% B to 80% B over 1 min. The composition was held at 80% B for 3 min, and then dropped to 2% B over 0.1 min before being held at 2% B for another 5 min. The 480 mass spectrometer was operated in a data-independent manner automatically switching between the acquisition of a single Orbitrap MS scan (350-1400 m/z and a resolution of 120k) and 50 MS2 scans (NCE 30%, 30k resolution, an Automatic Gain Control of 2000%, a mass range of 200-2000 m/z and a maximal injection time of 55 ms) of an isolation width of 13.7 m/z collected over the mass range of 361 to 1033 m/z.

#### DIA-based proteomic analysis

Proteomic datasets were searched using a DIA-library free approach using Spectronaut (version 17.6). For the analysis of host changes in response to *A. baumannii* intracellular growth, data files were searched against the *A. baumannii* Ab398 and Ab803 proteomes (NCBI accession number will be added) and the Mouse Proteome (Uniprot accession: UP000000058, downloaded 13/05/2023). To assess changes in intracellular *A. baumannii* proteome compared to static and LB growth, datasets were searched against either the *A. baumannii* Ab398 or Ab803 proteomes and the Mouse Proteome. For all searches oxidation of Methionine was allowed as a variable modification, Carbamidomethyl was set as a fixed modification of Cysteine, and protease specificity was set to Trypsin. Protein quantitation was undertaken using MaxLFQ (*95*) based analysis. For the analysis of changes within the host proteome, statistical analysis was undertaken using student t-tests within Perseus (v1.4.0.6) (*96*) with only proteins observed in at least three biological replicates of one group considered for analysis. Missing values were imputed based on the total observed protein intensities with a range of 0.3 σ and a downshift of 2.5 σ. Biological replicates were grouped together, and student t-tests were used to compare individual groups with a minimum fold change of +/- 1 considered for further analysis. Multiple hypothesis corrections were undertaken using a permutation-based FDR approach allowing an FDR of 5%. To allow the comparison of the proteome changes observed in *A. baumannii* Ab398 or Ab803 when grown within host cells to static and LB grown *A. baumannii* normalization was undertaken on the observed *A. baumannii* proteins within R with statistical analysis undertaken using Perseus. Only proteins observed in at least three biological replicates of one group were considered for analysis and missing values were imputed based on the total observed protein intensities with a range of 0.3 σ and a downshift of 2.5 σ. For statistical analysis biological replicates were grouped together, and student t-tests used to compare individual groups with a minimum fold change of +/- 1 considered for further analysis with the permutation-based FDR set to 5%. For host and bacterial comparisons data visualization was undertaken using ggplot2 in R (*97*).

### GSEA analysis

The gene set enrichment analysis (GSEA) was performed using the software GSEA (version 4.3.3). Imputed LFQ data were loaded into the software and pathway enrichment was conducted using the mouse molecular signature database MH: hallmark gene sets (mh.all.v2024.1). Enrichment scores were visualized using SRplot online platform (*98*).

### Bacterial homologs identification and COG annotation

Genes in two strains were considered homologs if they had at least 70% identity and 90% coverage. This analysis was performed using reciprocal BLAST hit (RBH) on the usegalaxy.eu platform. Bacterial protein fasta files were submitted to EggNOG to obtained predicted Cluster of Orthologous Groups (COG) annotation of each bacterial protein (*99*).

### Immunoblot analyses

Antibiotic protection assays were performed in J774.A1 as described above. At specified timepoints, cells were washed thrice with PBS and lysed with Laemmli loading buffer 2x (125 mM Tris -HCl, pH6.8, 4% SDS (w/v), 10% β-mercaptoethanol (v/v), 20% glycerin (v/v), 0.1% bromophenol blue (w/v). Samples were separated by SDS-PAGE, transferred to a nitrocellulose membrane, and proteins of interest were probed using polyclonal rabbit anti- IRG1 (1:2,000; gift from Pr. Anthony Orvedahl, Washington university in St Louis) and monoclonal mouse anti-β-actin (1: 5000; Sigma-Aldrich). IRDye-conjugated anti-mouse IgG and anti-rabbit IgG were used as secondary antibodies (1:10,000 for each; LI-COR Biosciences, Lincoln, NE), and blots were visualized with the Odyssey CLx imaging system (LI-COR Biosciences).

### Bacterial LOS visualization

Total membrane preparations were performed by cell lysis and ultracentrifugation. Cultures from mid-exponential phase were harvested by centrifugation at 6,500 rpm at 4°C for 10 minutes. The pellets were gently resuspended in PBS with complete EDTA-free protease inhibitor mixture (Roche Applied Science) followed by cell disruption. Centrifugation at 6,500 rpm at 4 °C for 5 minutes was performed to remove unbroken cells. Total membranes were collected by ultracentrifugation at 200,000 xg for 1 h at 4 °C and resuspended in PBS.

LOS content was measured according to Tsai and Frasch (*100*). Briefly, samples were diluted and treated with Proteinase K for 3h at 37°C prior to loading onto a 10-20% gradient SDS-PAGE gel. After running, gels were fixed overnight in 200 mL of 40% ethanol in 5% acetic acid. Next, gels were oxidized for 5 min in 100 mL of 0.7% fresh periodic acid in 40% ethanol and 5% acetic acid. Upon completion, gels underwent three washes (15 min each) in milliQ H_2_O. The gels were then stained for 10 min in the dark with 28 mL 0.1 M NaOH, 2 mL NH_4_OH, 5 mL 20% AgNO_3_, and 115 milliQ H_2_O. Gels underwent three additional washes prior to developing in 200 mL H_2_O with 10 mg citric acid and 100 μL formaldehyde.

### Immunofluorescence staining

1.3x10^5^ J774A.1 cells were seeded onto autoclaved glass coverslips in 24 well plates and incubated 16 to 18 h at 37 °C and 5% CO_2_. *A. baumannii* strains were prepared as per the antibiotic protection assay in PBS. Cells were infected with bacterial suspension to a final MOI=10. Plates were then centrifuged for 10 min at 200 xg to enhance bacterial contact with the host cells and incubated for 1 h at 37 °C and 5% CO_2_. Cells were washed thrice with PBS, and extracellular bacteria were killed by colistin treatment for 1 h. At the indicated time points, samples were fixed with 4% paraformaldehyde for 15 min at 37 °C and then stored in PBS until processing. Cells were incubated into a permeabilizing and blocking solution [PBS + 0.1% saponin + 0.5% BSA (Fisher BioReagents, BP9706100) + 10% hi-FBS (Corning)]. The glass coverslips were incubated with anti-*A. baumannii* primary antibodies produced in rabbit at a 1:100 dilution for 1 h at 37 °C (Antibody research corporation). Cells were then washed 3 times with washing solution (PBS + 0.1% saponin + 0.5% BSA) and incubated with the secondary antibody goat anti-rabbit: Alexa Fluor 488 (Invitrogen) at a 1:250 dilution, Alexa Fluor 555 phalloidin (0.33 μM; CST, #8953) and DAPI for 1 h at 37 °C. Samples were washed with PBS, rinsed with Milli-Q water, and the coverslip was mounted onto a slide using Invitrogen ProLong Gold Antifade Mountant (Invitrogen, P36930). Stained samples were analyzed by confocal microscopy.

### Confocal microscopy

Prepared slides were analyzed with a Zeiss LSM880 laser scanning confocal microscope (Carl Zeiss Inc.) equipped with 405nm diode, 488nm Argon, 543nm HeNe, and 633nm HeNe lasers. A Plan-Apochromat 63X (NA 1.4) DIC objective and ZEN black 2.1 SP3 software were used for image acquisition. Images were analyzed using ImageJ software (NIH, USA).

### Growth curves

*A. baumannii* strains were cultured overnight in M9 (1 g NH4Cl, 3 g KH2PO4, 6 g Na2HPO4, 4 g glucose, and 1 ml of 1 M MgSO4/L) with 0.1% casamino acids and 0.5, 5 or 20 mM of itaconic acid (Sigma). Alternatively, strains were cultured in LB broth, buffered LB pH5 (2-(*N*- morpholino-ethanesulfonic acid (MES) 100 mM) or in DMEM High Glucose supplemented with 10% hi-FBS at 37 °C under shaking conditions. Overnight bacterial cultures were centrifuged at 6500 rpm for 5 min and the pellets were washed twice with PBS. *A. baumannii* suspensions were diluted to OD_600nm_ = 0.01 in 150 μl of the media of interest in a 96-well round-bottom plates (Corning), covered with a Breatheasy film (Diversified Biotech) and incubated 16 h at 37 °C under shaking conditions. OD_600nm_ values were measured at 30 min intervals using a BioTek microplate spectrophotometer. The experiment was performed in biological triplicates with four technical replicates for each strain and experimental condition.

### Statistical analysis

For animal experiments, the two groups were compared using an unpaired t-test with Welch’s correction. For data from antibiotic protection assay, data from each timepoint was extracted, log-transformed, and a one-way ANOVA was performed with comparison of all samples to Ab398 wild-type. Statistical analyses were performed on Prism software.

## Data availability

Annotated Ab398 and Ab803 genomes were deposited on NCBI (bioproject number will be added here). The mass spectrometry proteomics data have been deposited to the ProteomeXchange Consortium via the PRIDE partner repository (ebi.ac.uk/pride/) with the dataset identifier PXD060577 (*101*).

## Figures

Prism software was used to display experimental data. The experimental workflows were made using bioRender (biorender.com) and Inkscape was used to create the working model and assemble all figures.

## Acknowledgement

This work was supported by funding to M.F.F. (R01AI166359 and R21AI181334). N.E.S was supported by an Australian Research Council (ARC) Future Fellowship (FT200100270), an ARC Discovery Project Grant (DP210100362), and a National Health and Medical Research Council Ideas grant (2018980). We thank the Melbourne Mass Spectrometry (MS) and Proteomics Facility of The Bio21 Molecular Science and Biotechnology Institute for access to MS instrumentation. We thank the imaging laboratory of the Molecular Microbiology Department at Washington University in St. Louis. We want to thank Wandy Beatty for her collaboration in obtaining the electron microscopy images shown in this work. We thank Anthony Orvedahl for the anti-IRG1 antibodies. The modern clinical isolate Ab803 (ABM452) used in this study was kindly provided by Mesoumeh Douraghi. We thank Dakota Hall for technical support and David Rosen and the members of the Feldman lab for critical reading of the manuscript. We thank Lance Bottini for his diligent care of our instruments.

## Author contributions

Conceived and designed the experiments: M.JM., G.DV. C.JL., N.E.S., and M.F.F. Performed the experiments: M.JM., C.JL., N.E.S. Analyzed the data: M.JM., N.E.S. Wrote the first draft of the manuscript: M.JM. All authors edited the manuscript.

## Competing interests

All authors declare that they have no competing interests.

## Data and materials availability

All data needed to evaluate the conclusions in the paper are present in the paper and/or the Supplementary Materials or have been deposited on public repositories. Additional data related to this paper may be requested from the authors.

## Notes

### Competing Interest Statement

The authors have declared no competing interest.

